# Notch signalling patterns retinal composition by regulating *atoh7* during post-embryonic growth

**DOI:** 10.1101/363010

**Authors:** Alicia Pérez Saturnino, Katharina Lust, Joachim Wittbrodt

## Abstract

Patterning of a continuously growing naive field in the context of a life-long growing organ, the teleost eye is of highest functional relevance. Intrinsic and extrinsic signals were proposed to regulate lineage specification in progenitors that exit the stem cell niche in the ciliary marginal zone (CMZ). The proper cell type composition arising from those progenitors is prerequisite for retinal function. Our findings in the teleost medaka (*Oryzias latipes*) uncover that the Notch–Atoh7 axis continuously patterns the CMZ. The complement of cell-types originating from the two juxtaposed progenitors marked by Notch or Atoh7 activity contains all constituents of a retinal column. Modulation of Notch signalling specifically in Atoh7-expressing cells demonstrates the crucial role of this axis in generating the correct cell type proportions. After transiently blocking Notch signalling, retinal patterning and differentiation is reinitiated *de novo*. Taken together we show that Notch activity in the CMZ continuously structures the growing retina by juxtaposing Notch and Atoh7 progenitors giving rise to distinct, complementary lineages, revealing a coupling of *de novo* patterning and cell-type specification in the respective lineages.

## Introduction

The central nervous system (CNS) presents an extraordinary diversity of neuronal cell types. Even now, the exact number of distinct neuronal cell types is unclear. Moreover, their lineage specification from a common progenitor pool is also very complex (Edlund and Jessell, 1999; Pearson and Doe, 2004). The retina, even though is part of the CNS, has a relatively simple cellular composition, which has been extensively studied (Bassett and Wallace, 2012). It consists of six neuronal cell types and one glial cell type, which are distributed into three nuclear layers: the outer nuclear layer (ONL) containing the rod and cone photoreceptors (PRC), the inner nuclear layer (INL) where bipolar cells (BC), amacrine cells (AC), horizontal cells (HC) and Müller glia (MG) cells are located and the ganglion cell layer (GCL) where the retinal ganglion cells (RGC) as well as some ACs reside. Its well-characterized structure together with its accessibility and easy manipulation put the retina in a privileged position to uncover the principles of lineage specification.

During lineage specification in retinal development, intrinsic as well as extrinsic factors influence the progression of progenitors through different competence states to achieve the production of the different cell types (Hufnagel and Brown, 2013; Livesey and Cepko, 2001). This process continues lifelong in constantly growing organisms such as fish and amphibians and is supported by stem cells, which reside in the most peripheral domain of the retina, the ciliary marginal zone (CMZ) (Easter and Johns, 1977; Hollyfield, 1968). Similarly to embryonic development, this pool of multipotent stem cells give rise to the whole spectrum of retinal cell types during post-embryonic growth (Centanin et al., 2011; Centanin et al., 2014). However, how lineage specification and patterning are coordinated in a continuously growing organ remains elusive.

Postembryonic growth shows some marked differences from embryonic development. Whereas new structures and organs need to be formed during embryonic development, already existing functional structures need to be expanded during post-embryonic growth. The Notch signalling pathway has been previously identified as extrinsic factor influencing cell fate decisions during retinal development (Andreazzoli, 2009; Livesey and Cepko, 2001). Despite the extensive studies in retinal development, little is known about cell specification of post-embryonic retinal stem and progenitor cells. The transmembrane receptor Notch as well as other components of the Notch signalling pathway have been reported to be expressed in the CMZ in frogs and fish suggesting its role in retinal post-embryonic growth (Dorsky et al., 1995; Raymond et al., 2006). However, this has not been addressed yet.

The potential implication of Notch signalling in cell fate specification during retinal post-embryonic growth is strongly supported by the classical role of Notch signalling in neural development: patterning regulation and cell fate determination (Louvi and Artavanis-Tsakonas, 2006). Notch signalling is known to regulate tissue diversification by generating a mosaic pattern: Notch signalling propagates in an equipotent tissue, generating a binary pattern, where adjacent cells differ from each other on the activation of the pathway (lateral inhibition). The activation of the pathway occurs when the transmembrane receptor Notch binds its ligand Delta, a protein located in the membrane of the neighbouring cell. This binding triggers proteolytic activity on the receptor releasing its intracellular domain, which translocates into the nucleus and regulates gene expression (Fiúza and Arias, 2007). Notch signalling downstream target genes, Hairy/E(spl)-related factors (Her factors in fish, Hes in mouse) function as transcriptional repressors (Borggrefe and Oswald, 2009). Among their targets, Her factors have been shown to act on basic helix loop helix (bhlh) proneural factors (Kageyama et al., 2007). Eventually, this results in a salt-and-pepper pattern of proneural gene expression (Lai, 2004; Neves et al., 2013). Amongst other bhlh factors, *atoh7* expression has been previously shown to be negatively regulated by Her factors during development (Maurer et al., 2014; Schneider et al., 2001). The role of Atoh7 in retinal development has been previously described in teleost fish. It has been shown to be necessary and sufficient for the development of RGCs (Kay et al., 2001). Atoh7-positive progenitors also give rise to ACs, HCs and PRCs during retinal development (Poggi et al., 2005). Interestingly, *atoh7* has been also shown to be expressed in the progenitor area of the post-embryonic teleost retina (Lust et al., 2016). However, the role for Notch signalling as well as its crosstalk with *atonal* genes in retinal post-embryonic growth is still unknown.

Here we show that Notch signalling is active in a subset of progenitors in the transit-amplifying zone of the CMZ in the Japanese rice fish medaka (*Oryzias latipes*). This progenitor population is fate-restricted to BCs, ACs and MG cells. Moreover, Notch signalling activation shows a mutually exclusive pattern with the expression of the basic helix loop helix transcription factor Atoh7. Manipulation of Notch signalling by targeted activation as well as its chemical inhibition demonstrate its crucial role in generating the correct cell type proportions within the Atoh7 lineage. All this occurs continuously *de novo* in the ciliary marginal zone where after transient Notch inhibition the Notch – atoh7 axis is reinitiated from scratch and maintained thereafter. Our data provide mechanistic insight into how a growing organ is patterned continuously de novo and how that this patterning in 2D, the juxtaposition of Notch and Atoh7 cells in the CMZ, impacts on the third dimension of cell type composition by distinct lineage specification.

## Results

### Notch signalling is active in a subset of retinal progenitors in the post-embryonic retina in medaka

Notch signalling is known to be active in Müller glia (MG) cells and the transit-amplifying zone of the CMZ in the zebrafish post-embryonic retina (Link and Darland, 2001; Raymond et al., 2006). Its role in MG cells, which are the retinal stem cells responsible for retinal regeneration in zebrafish, has been extensively studied (Wan and Goldman, 2017; Wan et al., 2012). However, the function of Notch signalling in lineage specification in the transit-amplifying zone of the CMZ is still unknown. We addressed this in the medaka retina. Here retinal stem cells residing in the CMZ have been recently characterised: they have been shown to be multipotent and the transcriptional network regulating their stemness has been also identified (Centanin et al., 2011; Centanin et al., 2014; Reinhardt et al., 2015).

To visualize active Notch signalling in the post-embryonic retina in medaka, the previously characterized *tp1-MmHbb::*d2GFP (*tp1::*d2GFP) Notch signalling reporter (Clark et al., 2012; Lust et al., 2016) was used. The reporter construct carries 6 copies of the *tp1* promoter, a Notch responsive promoter containing 2 RBP-Jk-binding sites, followed by a minimal promoter (mouse beta globin) and a destabilized GFP (d2GFP) (Fig. 1A). The *tp1*::d2GFP reporter line shows Notch signalling activation in various tissues such as thymus, brain and intestine in hatch medaka (8 days post fertilization at 28°C) (Fig. 1B). Notch signalling activation pattern is highly conserved and activation in these tissues had been previously reported in other organisms from flies to mouse (Bajoghli et al., 2009; Bigas and Espinosa, 2012; Bray, 2016; Siebel and Lendahl, 2017). In addition, we generated a second Notch reporter line for short-term lineage analysis: *tp1-MmHbb::*tagRFP (*tp1::*tagRFP). This line carries the same construct as the *tp1::*d2GFP line but the short live d2GFP is replaced by a tagRFP (Fig. 1C). TagRFP is not targeted to degradation and therefore has a long half-life allowing a short-term lineage tracing due to label retention (Merzlyak et al., 2007). The *tp1::*tagRFP reporter line shows activation in the same tissues as the *tp1::d2GFP* line, including the brain, the thymus and the intestine in a medaka hatchling (Fig. 1D).

**Figure 1.**
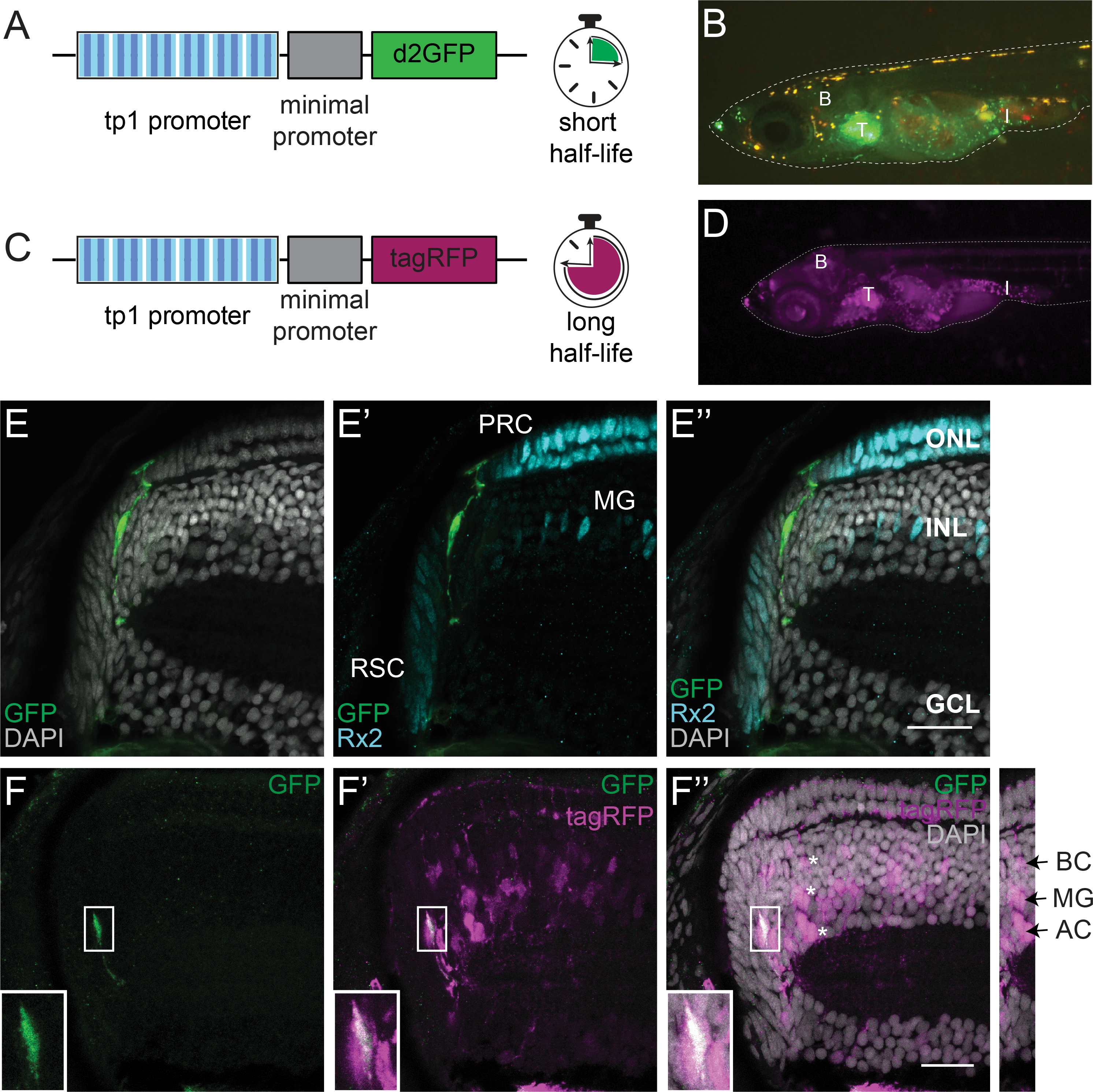
Notch signalling is active in a subset of retinal progenitors, which give rise to Müller glia cells, amacrine cells and bipolar cells during retinal post-embryonic growth in medaka. (A) The *tp1-MmHbb::*d2GFP (*tp1::*d2GFP) Notch signalling reporter contains 6 copies of the *tp1* promoter, a Notch responsive promoter (blue stripped boxes). Each *tp1* promoter contains 2 RBP-Jk-binding sites (dark blue stripes). The *tp1* promoter is followed by a minimal promoter (mouse β globin) and a destabilized GFP (d2GFP), which has a short half-life. (B) The *tp1::*d2GFP reporter line shows Notch signalling activation in various tissues such as brain (B), thymus (T) and intestine (I) in hatch medaka. (C) The *tp1-MmHbb::*tagRF (*tp1::*tagRFP) Notch signalling reporter contains the *tp1* Notch responsive promoter followed by a tagRFP, a very stable red fluorescent protein with a long half-life. (D) The *tp1::*tagRFP reporter line shows Notch signalling activation in hatch medaka in the same tissues as the previously described green reporter line: brain (B), thymus (T) and intestine (I). (E-E’’) Notch signalling (green) is active in a subset of retinal progenitors but it is neither active in RSCs nor in differentiated cells (PRCs and MG cells), here labelled with Rx2 (cyan). Nuclear labeling with DAPI is in grey (E’’). (F-F’’) The two Notch reporter lines (*tp1::*d2GFP in green and *tp1::*tagRFP in magenta) show overlapping activation only in the progenitor area (bottom left insets). Due to label retention, the *tp1::*tagRFP reporter line is also visible in differentiated cells (indicated by asterisks) derived from Notch-positive progenitors: Müller glia (MG) cells, amacrine cells (AC) and bipolar cells (BC). Scale bar is 20 μm. RSC: retinal stem cells; PRCs: photoreceptors; MG: Müller glia; ONL: outer nuclear layer; INL: inner nuclear layer; GCL: ganglion cell layer.

In the retina, we detected Notch signalling activation in a subset of progenitors (Fig. 1 E-E’’). Retinal progenitors are located in the transit-amplifying zone of the CMZ between the retinal stem cells (RSCs) and the central differentiated retina. RSCs are labelled by antibody staining against the transcription factor retinal homeobox gene two (*rx2*) (Reinhardt et al., 2015). They are located at the most periphery of the retina. The differentiated retina occupies the layered and most central part of the tissue. Among the other retinal cell types, it contains PRCs and MG, which also express *rx2*. Importantly, Notch signalling was not detected in RSCs. MG cells and PRCs do not show Notch signalling either. This is in contrast to zebrafish in which Notch signalling is active in MG cells (Wan and Goldman, 2017). The non-overlap of Notch signalling with either RSCs or MG cells or PRCs can be clearly appreciated in the *tp1::*d2GFP retina 3D reconstruction from a frontal (Fig. S1A,A’) and lateral (Fig.S1 B,B’) view. Our data show that Notch signalling is confined to a subset of progenitors in the CMZ of the medaka retina.

### Notch-positive progenitors give rise to MG, AC and BC

Notch signalling has been previously shown to be involved in cell-fate choices in the embryonic vertebrate retina (Andreazzoli, 2009; Livesey and Cepko, 2001). During post-embryonic retinal growth we hypothesize lineage specification to occur in the transit-amplifying zone of the CMZ, between RSCs and the more centrally located terminally differentiated retina. Factors involved in lineage specification during retinal development such as *atoh7* and *neuroD* have been shown to be expressed in that zone (Lust et al., 2016; Poggi et al., 2005; Raymond et al., 2006; Taylor et al., 2015). Consistently, the activation of Notch signalling in the transit-amplifying zone of the CMZ points towards a role for Notch signalling in cell fate specification during post-embryonic growth. Thus, we next addressed the differentiation potential of the Notch-positive progenitor pool.

To assess that we crossed the *tp1::*tagRFP reporter line to the *tp1::*d2GFP reporter. In the retina of these double transgenic fish, we distinguished two cell populations. One population located towards the periphery of the CMZ was double positive reflecting actual Notch signalling activation (Fig. 1F-F’’, insets). The second population located more centrally in the differentiated retina was only positive for tagRFP and corresponds to the cells that derived from a Notch-positive progenitor pool (Fig. 1F-F’’). Interestingly, these cells were only located in the INL and were identified as BCs, MG cells and ACs based on morphology and position: BCs are located in the apical part of the INL; the nucleus of MG cells is elongated and located in the centre of the INL; ACs have a round nucleus and are located in the basal part of the INL (Fig. 1F’’ asterisks and inset on the right). No tagRFP-positive nucleus was observed in the ONL or GCL. The tagRFP signal that can be observed in those layers corresponds to cellular projections of BCs and MG cells. In addition, we validated the identity of the descendants of Notch-positive progenitor cells by co-staining with cell type specific markers (Fig. S2). Taken together these results show that the lineage of the Notch-positive progenitor pool comprises a subset of retinal cell types, belonging to the INL.

### Notch signalling activation and *atoh7* expression show mutually exclusive patterns in the progenitor area of the post-embryonic medaka retina

Notch-positive progenitors are committed to BCs, MG cells and ACs. These progenitors comprise only a subset of progenitors in the CMZ and do not generate the complete spectrum of retinal cells types, Therefore, another pool of progenitors must give rise to RGCs, PRCs and HCs, complementing the Notch lineage.

The bhlh transcription factor Atoh7 is well known for its role during retinal development in vertebrates (Kay et al., 2001; Ohnuma et al., 2002). *atoh7* is expressed in the final divisions of retinal progenitors and known to be necessary for their differentiation into RGCs. The lineage of Atoh7-positive retinal embryonic progenitors comprises RGCs, PRCs, ACs and HCs (Poggi et al. 2005). It has been recently shown that *atoh7* expression is not restricted to embryonic development but a subset of progenitors in the CMZ also expresses *atoh7* during post-embryonic growth. This pool of progenitors has the same potential as its embryonic counterpart (Lust et al., 2016).

In order to investigate how Atoh7-positive progenitors localize within the CMZ with respect to the Notch-positive progenitors, the *atoh7::*GFP reporter line (Lust et al., 2016) was crossed to the *tp1::*tagRFP reporter line (Fig. 2A). In double transgenic fish, Notch- and Atoh7-positive progenitors showed a mutually exclusive pattern in the progenitor area of the CMZ: in that area of the retina, the progenitor cells are either Notch-positive or Atoh7-positive but never positive for both of them (Fig 2B-C’’). This mutually exclusive arrangement of Notch-positive and Atoh7-positive cells is resembling the Notch-Delta lateral inhibition pattern previously described in numerous organisms and tissues based on RNA expression patterns (Chitnis et al., 1995; Kiernan et al., 2005; Parks et al., 1997). Interestingly, a third population of progenitors negative for both, Notch and Atoh7 was also observed. These results indicate that Notch signalling and *atoh7* expression are tightly coordinated to form a mutually exclusive pattern. Moreover, the lineages derived from Notch- and Atoh7-postive progenitors show a striking complementarity. This let us to hypothesise a crosstalk between Notch signalling and *atoh7* expression to regulate cell fate and lineage restriction during post-embryonic growth.

**Figure 2.**
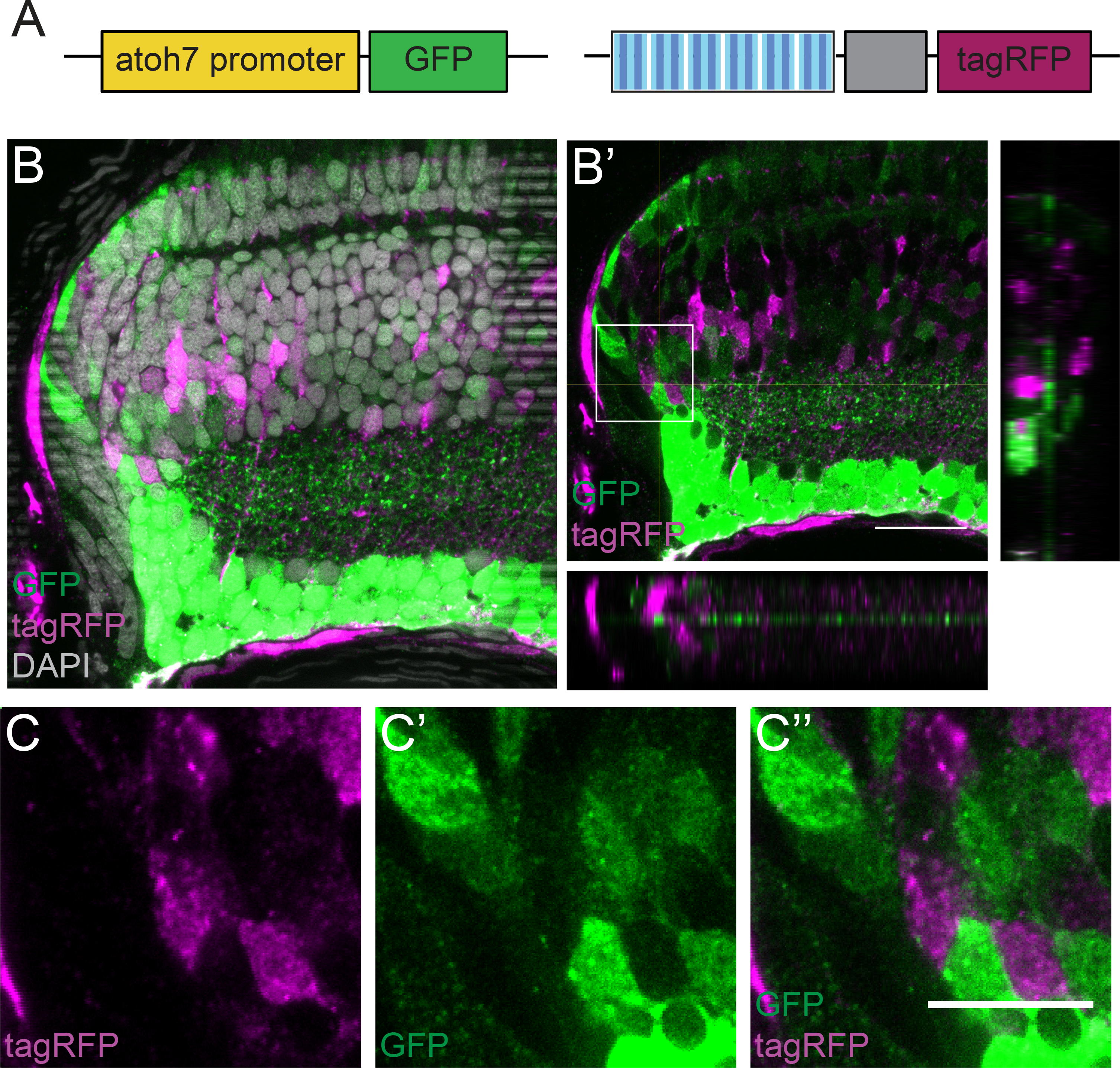
Notch signalling activation and *atoh7* expression show a mutually exclusive pattern in the progenitor area of the post-embryonic medaka retina. (A) The *atoh7* reporter line containing the *atoh7* promoter followed by a GFP was crossed to the *tp1::*tagRFP Notch reporter. (B,B’) Notch signalling activation (magenta) and *atoh7* expression (green) show mutually exclusive patterns in the progenitor area of the CMZ. DAPI is shown in grey in B. Orthogonal views (XZ, YZ) of the progenitor area in B’. Scale bar is 20 μm. (C-C’’) Higher magnification of the progenitor area indicated with a square in B’. TagRFP is shown in magenta (C), GFP in green (C’) and the merge is shown in C’’. n=8 fish. Scale bar is 10 μm.

### Toolbox for targeted activation of Notch signalling in Atoh7-positive progenitors

To address the role of the Notch-Atoh7 crosstalk in cell fate specification in the retina, we intended to interfere with the tightly regulated balance in the progenitor area by activating Notch signalling in Atoh7-positive progenitors and analysing the effects on lineage specification.

To achieve this, we generated an *atoh7::*^ERT2^ Cre line as a recombination driver (Fig. S3). This line was validated for recombination in the *atoh7* expression domain in the CMZ. For that we crossed this driver line with a control line carrying a construct that results in a colour switch (from mCherry to H2B-GFP) upon recombination (GaudíRSG line, Centanin et al. 2014) (Fig. 3A). Immediately after recombination, we observed GFP-positive cells in the *atoh7* expression domain in the CMZ indicating specific recombination in Atoh7-progenitors (Lust et al., 2016) (Fig. S3A-A’’). We validated the specific activity of the *atoh7::*^ERT2^ Cre in progenitor cells by lineage analysis. One-month after recombination of GaudíRSG by induction of *atoh7::*^ERT2^ Cre, we detected terminating clones in a ring around the embryonic retina (Fig. S3B-B’’) identifying the initially targeted Atoh7-positive cells as progenitors with limited proliferative potential.

**Figure 3.**
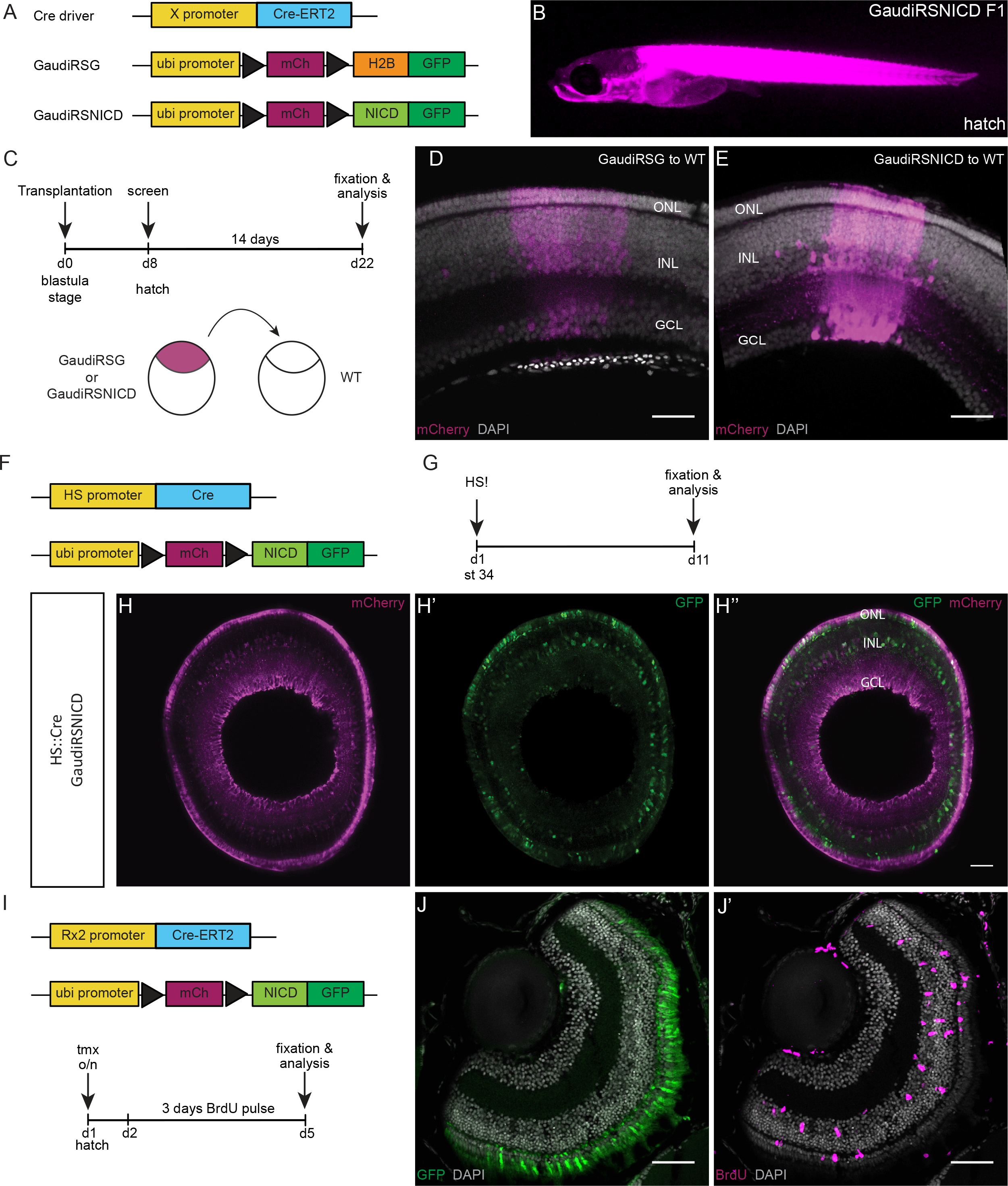
The GaudíRSNICD line recombines ubiquitously and is functional. (A) Specific promoters (X promoter) can be used to drive expression of an inducible Cre recombinase in specific cell populations to achieve targeted recombination of the GaudíRSG construct and the GaudíRSNICD. The GaudíRSG construct is the control construct: it switches from mCherry to H2B-GFP upon recombination. Black triangles represent LoxP sites. The GaudíRSNICD construct is the experimental construct: it switches from mCherry to Notch intracellular domain (NICD) fused to a GFP upon recombination. (B) GaudíRSNICD fish after hatching showing ubiquitous mCherry expression (magenta). (C) Transplantations were performed at blastula stage (day 0, d0) from GaudíRSG or GaudíRSNICD embryos into WT embryos (see scheme below). The fish were screened for mCherry expression at hatch (d8) and grown for 14 days. After that, they were fixed and analysed (d22). (D, E) The transplanted cells from both lines (GaudíRSG in D and GaudíRSNICD in E) integrated in the host retina and differentiated into all cell types (Centanin et al., 2011). mCherry-positive cells are shown in magenta and nuclear labelling with DAPI in grey. The retinal cellular layers are also indicated: outer nuclear layer (ONL), inner nuclear layer (INL) and ganglion cell layer (GCL). Scale bar is 40 µm. (F) GaudíRSNICD fish were cross to a heatshock (HS)::Cre line. (G) The double positive fish were heatshocked (HS!) at stage 34 (st. 34) and fixed and analysed 10 days later. (H-H’’) Recombined retinae show mCherry expression (magenta, H) and recombination in all three cellular layers (GFP-positive cells, green, H’). H’’ shows the overlay of H and H’. The three retinal layers are indicated: ONL, INL, GCL. Scale bar is 40 µm. (I) Recombination of the GaudíRSNICD construct was induced with an inducible Cre recombinase under the control of *rx2* promoter. After overnight tamoxifen induction, the fish were incubated for 3 days in BrdU and then fixed and analysed. (J) Recombination in *rx2*-expressing cells such as PRCs could be observed. (J’) Massive proliferation in the central retina was detected by immunostaining against BrdU (magenta) (n = 7 fish). Scale bar is 40 µm. This result recapitulated the previously reported phenotype observed upon Notch activation in Rx2-ositive cells (Lust et al., 2016).

To specifically activate Notch signalling in Atoh7-positive cells, we combined the *atoh7*::^ERT2^ Cre driver line with an experimental construct switching from ubiquitous mCherry to GFP fused to the Notch intracellular domain (GaudíRSNICD line, Fig. 3A). Whereas recombination of the control GaudíRSG construct only induces a colour switch, recombination of the GaudíRSNICD construct will initiate activation of the Notch signalling pathway in combination with a colour readout. Since the recombination occurs at the genetic level, the switch will result in a permanent labelling, which allows following the lineage in both conditions.

The GaudíRSNICD line was first validated for ubiquitous expression, recombination and functionality. Hatchling fish from this line show ubiquitous mCherry expression (Fig. 3B). To further confirm the ubiquitous expression of the construct at cellular level in the retina, we performed transplantations at blastula stage from GaudíRSNICD into WT fish (Fig. 3C). As control for being ubiquitous we used the previously characterized line GaudíRSG (Centanin et al., 2014). The transplanted fish were screened at hatching stage for red fluorescence in the eye and grown for two weeks. After that period of time, they were fixed and stained for mCherry (Fig. 3C). Both, GaudíRSG and GaudíRSNICD, generated Arched Continuous Stripes (ArCoSs, Centanin et al. 2014) that show mCherry signal in all three cellular layers (ONL, INL, GCL) and therefore in all retinal cell types (Fig. 3D,E). These results confirm that the line expresses ubiquitously and the construct is actively transcribed in all retinal cell types.

Secondly, the line was validated for recombination. Recombination was induced with Cre recombinase under the control of the heatshock promoter (HS) at embryonic stage 34 (st. 34) (Fig. 3F). The fish were subsequently fixed and analysed 10 days after recombination (Fig. 3.G). The analysed retinae showed recombination in all three cellular layers of the retina (Fig. 3H-H’’).

Finally, the line was validated for being functional. To that end, the GaudíRSNICD line was crossed to the *rx2*:: ^ERT2^ Cre line. Notch signalling activation in Rx2-positive cells triggers massive proliferation in the central differentiated retina (Lust et al., 2016). We use that effect to validate the GaudíRSNICD line. We crossed it to the *rx2*::^ERT2^ Cre line and reproduced the previously reported effects (Fig. 3I-J’). Upon recombination, GFP can be observed in Rx2-positive cells (Fig. 3J). The massive proliferation observed in the central retina exactly reflects the previously described phenotype (Fig. 3J’). All together, these results show that the GaudíRSNICD line can be ubiquitously activated to full functionality.

### Targeted activation of Notch in Atoh7-positive progenitors shifts their potential

After validation of the *atoh7*::^ERT2^ Cre and the GaudíRSNICD lines, we addressed the impact of the Notch-Atoh7 crosstalk on the specification of the resulting lineages. To that end, *atoh7*:: ^ERT2^Cre fish were crossed to GaudíRSNICD or to GaudíRSG controls (Fig. 4A). Cre recombinase was activated at hatching stage by an overnight tamoxifen induction. The subsequent application of BrdU was used to leave a ‘timestamp’ highlighting the cells actively dividing at the time point of recombination. After a growth time of two weeks, the lineage was analysed by counting GFP-positive cells in each cellular layer located peripheral to the BrdU ‘timestamp’ (Fig. 4B).

**Figure 4.**
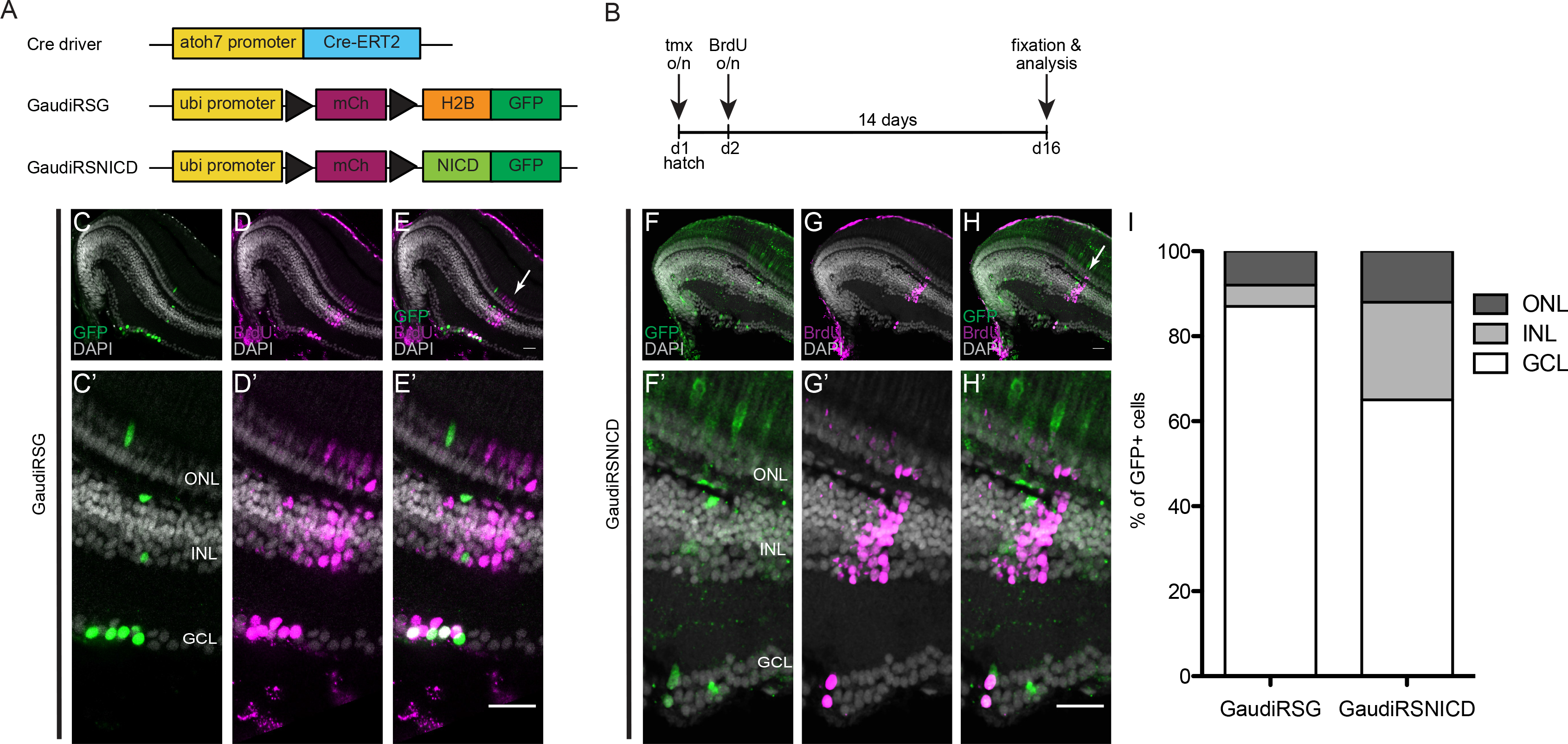
Notch signalling activation in Atoh7-positive progenitors shifts cell fate ratios during post-embryonic growth. (A) The *atoh7* promoter was used to drive an inducible Cre recombinase exclusively in Atoh7-positive cells to achieve targeted recombination of the GaudíRSG construct (control construct: switch from red to green upon recombination) and the GaudíRSNICD construct (functional construct: switch from red to Notch intracellular domain with a GFP upon recombination). (B) Recombination was triggered with an overnight induction with tamoxifen at hatch. The fish were incubated in BrdU overnight to label the induction time point and grown for 14 days afterwards. At day 16 the fish were fixed and the lineage was analysed. (C-H’) GaudíRSG fish as well as GaudíRSNICD fish show recombined cells (GFP-positive, green) belonging to the Atoh7 lineage. These recombined cells are located at the BrdU (magenta) stripe or more peripheral. Scale bar in E, E’, H and H’ is 20 μm. (I) Stacked column graph showing the percentage of cells in each layer derived from Atoh7-positive progenitor in control (GaudíRSG) and experimental (GaudíRSNICD) conditions. GCL; ganglion cell layer, INL; inner nuclear layer, ONL; outer nuclear layer.

Induction in the GaudíRSG control line confirmed the previously described lineage: Atoh7-positive cells give rise to RGCs, ACs, HCs and PRCs (Fig. 4C-E’). Upon Notch activation, qualitatively the same cell type composition was observed in the GaudíRSNICD line (Fig. 4F-H’). However, the proportions of cell types generated in each condition were strikingly different (Fig. 4I). We observed a severe loss of RGCs (to 74%) in response to Notch activation in Atoh7-positive cells that was compensated by a 5-fold gain in cell numbers in the INL (Fig. 4I, Fig. S4A,B). The generation of PRCs was not significantly altered by the activation of Notch signalling in Atoh7-positive progenitors (Fig. 4I, Fig. S4C). These results indicate that activation of Notch signalling shifts the proportions of cell types generated from the Atoh7-positive progenitors while it does not impact on their differentiation potential suggesting a regulation of Athoh7 by Notch signalling. Thus, continuous activation of Notch signalling shifts cell fate ratios within the Atoh7 lineage and leads to an increase in cells in the INL at the expense of RGCs.

### Inhibition of Notch signalling increases the number of Atoh7-positive progenitors

Regulation of *atoh7* expression by Notch signalling had been previously reported in the developing mouse retina (Maurer et al., 2014). To address whether Notch signalling regulates the expression of *atoh7* in the continuously growing medaka retina we assessed the expression of *atoh7* upon Notch inhibition. To do so we employed the Notch inhibitor LY411575 which had been successfully used to block Notch signalling in zebrafish (Mizutari et al., 2013; Romero-Carvajal et al., 2015). We tested the drug on hatching stage medaka and validated its specificity for the Notch signalling pathway in *tp1::*d2GFP reporter animals. A complete Notch signalling inhibition was achieved after 4 days of treatment (Fig. S5). After the validation of the inhibitor we made use of double reporter fish: *tp1::*d2GFP and *atoh7::*lyntdTomato (Fig. 5A). We incubated these double reporter fish at hatchling stage with DMSO or Notch inhibitor for 4 days and fixed immediately after. These fish were analysed by immuno-staining against the respective fluorescent proteins (Fig. 5B). Upon Notch inhibition, the number of Atoh7-positive cells in the progenitor area of the CMZ was almost doubled and the domain is more apically expanded (Fig. 5C-E’’). This shows a repression of *atoh7* expression by Notch signalling in the post-embryonic medaka retina. Inhibition of Notch signalling de-represses *atoh7* expression and consequently more cells express *atoh7*, indicating the regulation of *atoh7* expression by Notch signalling.

**Figure 5.**
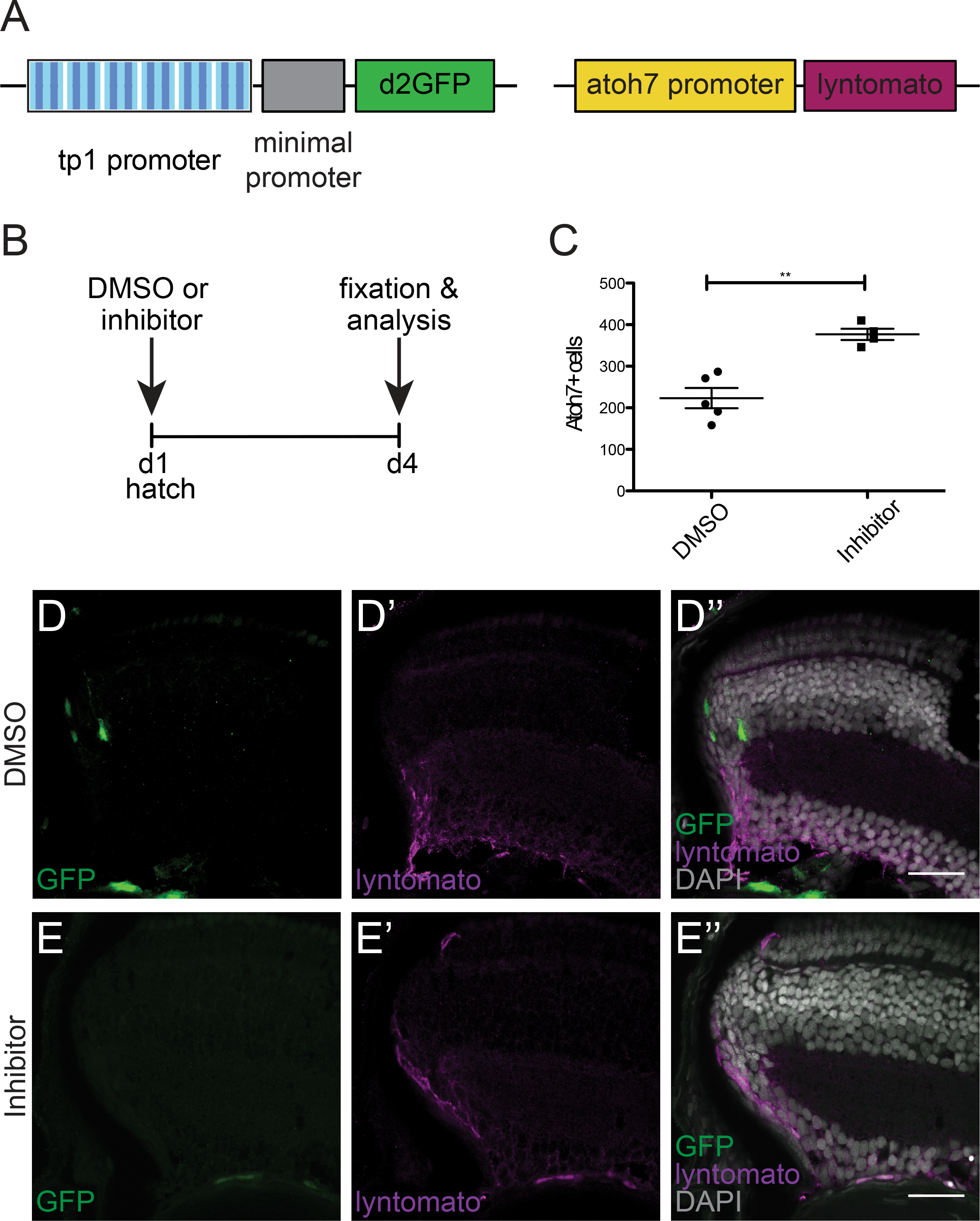
Notch signalling inhibition increases the number of Atoh7-positive progenitors. (A) Fish at hatching stage from a double reporter line carrying the *tp1::*d2GFP reporter and the *atoh7::lyntdTomato* reporter were used for this experiment. (B) The fish were treated with BrdU and DMSO (control) or Notch inhibitor (LY411575) for 4 days. At day 4, the fish were fixed and analysed. (C) Quantification of Atoh7-positive cells shows an increase upon Notch signalling inhibition. **p=0,0014 (1116 cells in 5 control fish and 1507 cells in 4 inhibited fish). (D-D’’) Control fish show Notch signalling activation (GFP, green) and a restricted domain of Atoh7-positive cells (lyntdTomato, magenta). (E-E’’) Notch inhibited fish do not show Notch signalling activation and the *atoh7* expression domain is expanded. Scale bar in D’’ and E’’ is 20 μm.

### Notch inhibition blocks PRC generation and increases the number of cells in the INL

We next addressed the impact of Notch inhibition on the Atoh7 lineage. We treated wild type fish at hatching stage for 4 days with either the LY411575 inhibitor or DMSO as control and applied BrdU as timestamp. After 4 days, the treatment was stopped and the animals were placed back in normal fish water for two weeks. Afterwards the fish were fixed and the retinal cell type composition analysed (Fig. 6A).

**Figure 6.**
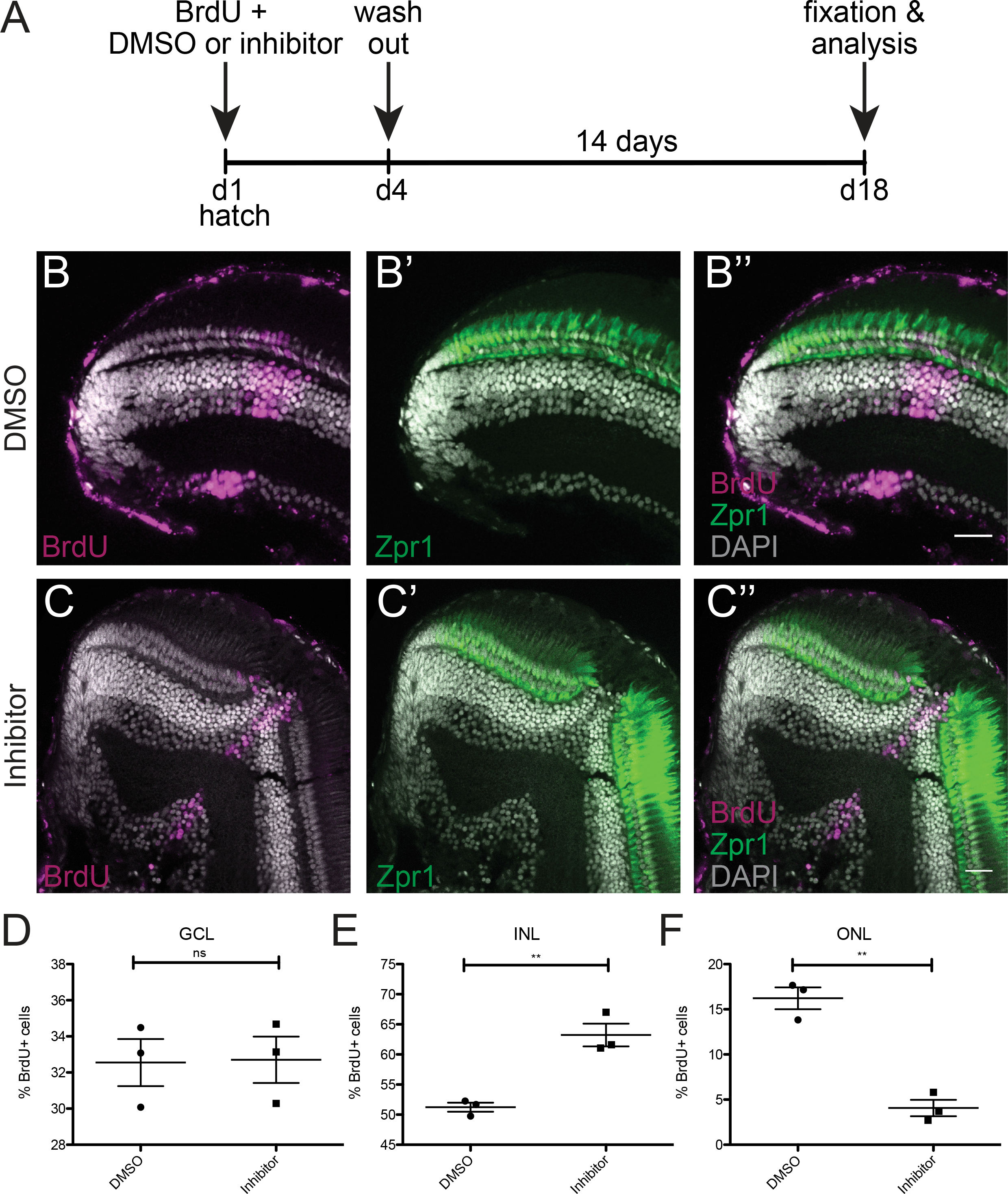
Notch signalling inhibition shifts cell fate ratios within the Atoh7 lineage. (A) Wild-type fish were treated at hatch with BrdU and DMSO (control) or Notch inhibitor (LY411575) for 4 days. At day 4 the BrdU and the DMSO or the inhibitor were washed out and the fish were grown for 14 days. At day 18, the fish were fixed and analysed. (B-B’’) Control retinae display a normal morphology and the BrdU (magenta) stripe corresponding to the treatment time contains cells in the three cellular layers. The PRCs labelled with Zpr-1 (green) also display a normal morphology. (C-C’’) Treated retinae display a disrupted morphology at the BrdU stripe, lacking the PRCs. (D-F) Quantification of BrdU-positive cells located in each layer in control conditions as well as in Notch-inhibited conditions reveals an increase in cells added to the INL and a decrease in photoreceptors located in the ONL. (D) GCL:^ns^ p=0.9372 (710 cells in 3 control retinae and 1303 cells in 3 inhibited retinae); (E) INL: **p=0.0042 (1104 cells in 3 control retinae and 2443 cells in 3 inhibited retinae); (F) ONL: **p=0,0013 (334 cells in 3 control retinae and 169 cells in 3 inhibited retinae). Scale bar is 20 μm.

Control fish were unaffected and before and after the BrdU timestamp showed a morphologically normal retina with the three classical retinal layers (Fig. 6B-B’’). In contrast, the Notch-inhibited retinae showed a strong response to the absence of Notch signalling and the subsequent *atoh7* upregulation (Fig. 6C-C’’). In the treated domain marked by the BrdU timestamp, the retina lacks the PRCs in the ONL as shown by absence of the cone photoreceptor marker Zpr-1. Accordingly, Notch inhibition results in an increase in the cell numbers composing the INL. Quantification of BrdU-positive cells located in each layer in each condition showed a 4-fold decrease in cells of the ONL (PRCs absent) and a 123% increase in cells of the INL as a result of Notch inhibition (Fig. 6D-F).

Together with the results in response to the targeted activation of Notch, our data indicate that Notch signalling is necessary for achieving the proper cell type proportions within the Atoh7 lineage. The fact that retinal growth and differentiation is properly reinstated after the extended and complete inhibition of Notch signalling indicates that the Atoh7-Notch pattern is re-established *de novo* and is crucial for the semi-crystalline architecture of the retina.

## Discussion

Notch signalling is known to strongly impact on neural development influencing multiple aspects of this complex process (Louvi and Artavanis-Tsakonas, 2006). Several studies had also addressed Notch in the developing retina, as part of the nervous system (Del Bene et al., 2008; Perron and Harris, 2000). However, even though expression of Notch pathway components has been reported in the CMZ of lifelong growing organisms such as *Xenopus* and zebrafish (Dorsky et al., 1997; Raymond et al., 2006) its role in the continuous establishment of regularly patterned 3D neural columns of retinal cell types during the continuous growth in the post-embryonic retina had not been addressed so far. Here we employ Notch pathway sensors to show that Notch signalling is active in a regularly spaced subset of progenitors in the transit-amplifying zone of the CMZ in medaka. This population is fate-restricted to cells exclusively located in the INL: BCs, ACs and MG cells. Notch signalling is active in a pattern that is mutually exclusive with the expression of the transcription factor Atoh7 in the transit-amplifying zone of the CMZ. Those Atoh7-positive progenitors are fated to become RGCs, HCs, ACs and PRCs (Lust et al., 2016), showing a striking complementarity to the Notch lineage itself. Modulation and manipulation of Notch signalling by genetically targeted activation of the pathway in Atoh7-positive cells as well as by chemical interference demonstrate the crucial role for Notch signalling in generating the correct cell type distribution and proportion within the Atoh7 lineage.

The immediate impact of Notch signalling on *atoh7* expression in neighbouring cells in the post-embryonic retina and the resulting mutually exclusive pattern of expression/activity collectively argue for a lateral inhibition scenario. We propose that the Notch – Atoh7 pattern in the transit-amplifying zone of the CMZ is shaping retinal architecture during the continuous post-embryonic growth (Fig. 7): adjacent cells show an opposite status regarding Notch signalling activity. Active Notch signalling in one cell represses *atoh7* expression and fixes a range of cell types that Notch positive progenitors give rise to (Notch lineage). Together with the descendants of the neighbouring Atoh7-positive cell (Atoh7 lineage) both establish the full complement of retinal cell types resulting in functional retinal columns in 3D. Strikingly, ACs are present in both lineages consistent with the multiple subtypes of ACs that have been identified, suggesting that each lineage is giving rise to different subtypes (Jusuf et al., 2011; Jusuf et al., 2012). Although the combination of both lineages covers all major retinal cell types, combining the two reporter lines carrying stable fluorophores, we observed cells in the central retina, which were negative for both reporters. It is known that not only ACs but also several other retinal cell types can be further defined into subtypes (Jusuf et al., 2012; Kim et al., 2008; Schmidt et al., 2011; Suzuki et al., 2013; Wan and Goldman, 2017). Or data indicate retinal subtypes that are neither generated from Atoh7-positive nor from Notch-positive progenitors. The presence of a third population of retinal progenitors in the CMZ, which was also negative for both Notch signalling and *atoh7* expression, supports the hypothesis of an additional mechanism to achieve the full spectrum of cellular diversity of the retina during post-embryonic growth further extending the complexity of the lateral inhibition scenario.

**Figure 7.**
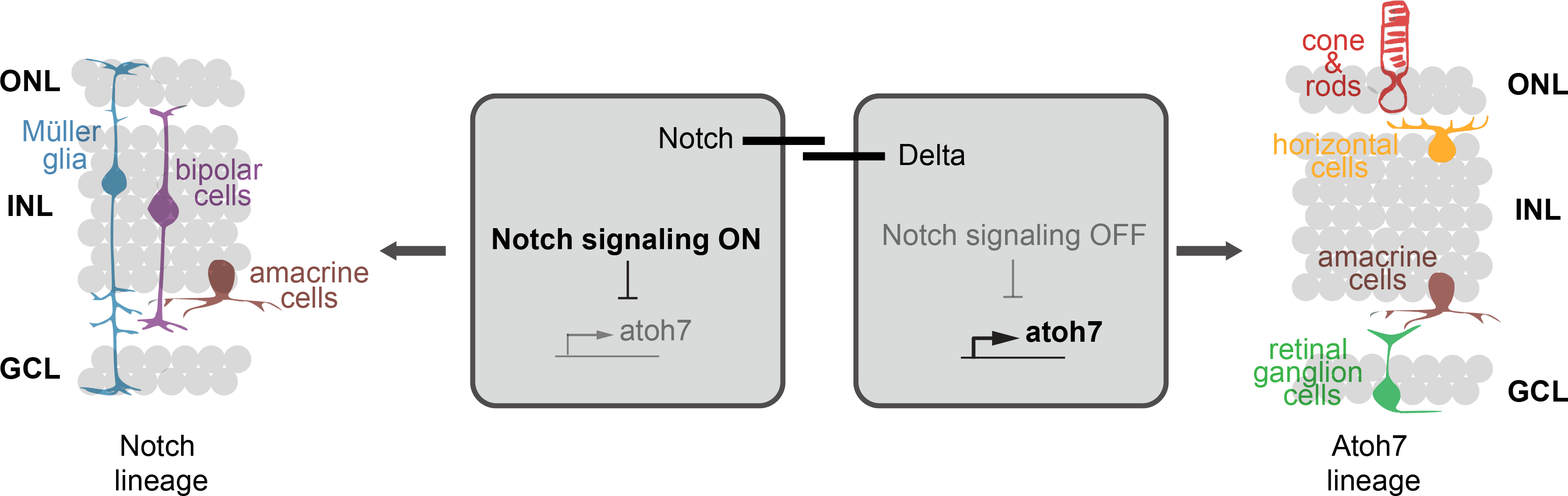
Notch signalling regulates the expression of *atoh7* by lateral inhibition. The cells that exhibit Notch signalling activation will repress *atoh7* expression and give rise to the cell types comprised in the Notch lineage, namely MG cells, BCs and ACs (right). In the neighboring cell, Notch signalling will be inactive and therefore *atoh7* can be expressed. Those cells will give rise to cell types comprised in the Atoh7 lineage: PRCs, ACs, HCs and RGCs (left). Scheme adapted from (Centanin et al., 2014).

Breaking the balance in the lateral inhibition scenario by activating or inhibiting Notch signalling impacts on the Atoh7 lineage indicating a crosstalk between Notch and Atoh7. Our results show that this crosstalk is accomplished by the repression of *atoh7* expression by Notch signalling. A direct repressive regulation of *atonal* genes by the Notch downstream factors (Her) has been previously shown in different model organisms and several tissues such as the inner ear, the lateral line and the intestine (Matsuda and Chitnis, 2010; Neves et al., 2013; VanDussen et al., 2012; Wilson et al., 2016). This regulation is a very conserved mechanism to segregate lineages. In the inner ear of chicken, mouse and zebrafish, Notch signalling specifies the non-sensory lineage, which includes all support cells. In the neighbouring cells, Notch signalling is blocked by lateral inhibition and here *atoh1* expression specifies hair cell fate (Atkinson et al., 2015; Kiernan, 2013; Neves et al., 2013). Similarly, in the murine intestine, for example, Hes1 has been shown to promote the absorptive lineage by repressing *atoh1* expression (Fre et al., 2005; Jensen et al., 2000). *atoh1* is then expressed in the other intestinal lineage, the secretory lineage (Koch et al., 2013; VanDussen et al., 2012). Taken together, the Notch – *atonal* axis is a recurrently employed mechanism to segregate complementary lineages in complex 3D tissues.

In the embryonic retina, a regulation of proneural *atonal* genes by Notch signalling had been proposed in multiple model organisms based on modelling results and *in situ* analysis. Our results experimentally substantiate this proposed regulation of *atonal* genes by Notch signalling. We show that Notch signalling modulations immediately impact on the Atoh7 lineage via influencing *atoh7* expression. Modelling *Drosophila* eye development, it has been proposed that active Notch signalling inhibits R8 photoreceptor fate by inhibiting *atonal* genes within the same cell. In contrast, in the neighbouring cell, where Notch signalling is laterally inhibited, *atonal* genes can be expressed allowing differentiation into R8 (Baker et al., 1996; Graham et al., 2010). In chicken, Notch signalling has been shown to inhibit RGC fate arguing for an influence of Notch signalling on the Atoh7 lineage (Austin et al., 1995). Similarly, manipulations on Notch signalling in the developing mouse retina affect the cell types belonging to the Atoh7 lineage. With an inducible deletion of the Notch downstream effector RBPJk, it was shown that autonomous RBPJk inhibits RGC and photoreceptor fates (Riesenberg et al., 2009). Another study further substantiated a regulation of *atoh7* expression by Notch signalling in the developing mouse retina (Maurer et al., 2014). Our direct modulation of Notch signalling and its impact on *atoh7* in conjunction with those data demonstrate that the Notch – *atonal* axis is fundamental for the segregation of retinal lineages to ultimately achieve the correct cell type composition and pseudo-crystalline architecture of the retina.

To address the link between growth and continuous patterning of the differentiating retina it was crucial to address whether the pattern established by the Notch – *atonal* axis is perpetuated from the already differentiated central retina and thus impacts on the newly forming retinal columns or whether it is *de novo* established in a continuous fashion in the retinal progenitor cells. Our experiments completely blocking Notch signalling were disconnecting the existing, highly patterned central retina form the newly forming tissue originating from the distal CMZ. Strikingly, even though cells are disturbed in their fate impacting on retinal lamination in the domain experiencing the Notch signalling block, the retina continues growing and re-instates proper lamination and differentiation *de novo*. This uncovers an effective self-organization capacity establishing retinal patterning in the growth zone of the fish retina with striking parallels to the proposed self-organisation driven by Notch-Delta lateral inhibition interactions in the sensory organ in *Drosophila* (Corson et al., 2017). In these diverse contexts, the Notch – *atonal* axis acts a fundamental and evolutionarily highly conserved mechanism combining pattern establishment and cell lineage specification to shift the temporal specification axis into the third dimension of cell types arranged in the retinal column.

## Acknowledgements

We thank C. Helker and D. Stainier for *tp1*::tagRFP reporter construct, T. Tavhelidse and B. Wittbrodt for designing and establishing the *atoh7*::^ERT2^ Cre construct and R. Sinn for the *atoh7*::lyndTomato medaka line. We are grateful to A Saraceno, E Leist and M Majewski for fish husbandry. We thank L. Centanin and all the members of the Wittbrodt group for constructive feedback on the project and critical reading of the manuscript. A P-S and KL were members of HBIGS, the Heidelberg Bioscience International Graduate and were both supported by a LGFG Fellowship. This work was supported by the European Research Council (GA 294354-ManISteC to JW) and the German Science Foundation (DFG, SFB 873 TP A3 to JW).

## Material and methods

### Animals and transgenic lines

Medaka (*Oryzias latipes*) used in this study were kept as closed stocks in accordance to Tierschutzgesetz 111, Abs. 1, Nr. 1 and with European Union animal welfare guidelines. Fish were maintained in a constant recirculating system at 28°C on a 14 h light/10 h dark cycle (Tierschutzgesetz 111, Abs. 1, Nr. 1, Haltungserlaubnis AZ35– 9185.64 and AZ35–9185.64/BH KIT). The following stocks and transgenic lines were used: wild-type Cabs, GaudíRSG (Reinhardt et al., 2015), *GaudíRSNICD*, rx2::^LoxP^ N3ICD (Lust et al., 2016), *atoh7*::iCre, *rx2*::iCre (Reinhardt et al., 2015), *HS::Cre* (Centanin et al., 2014) *tp1-MmHbb*::d2GFP (Lust et al., 2016), *tp1-MmHbb*::tagRFP, *atoh7*::GFP (Lust et al., 2016), *atoh7*::lyndtomato. All transgenic lines were created by microinjection with Meganuclease (I-SceI) in medaka embryos at the one-cell stage, as previously described (Thermes et al., 2002), except for *tp1-MmHbb*::tagRFP, which was created by microinjection with Tol2.

### BrdU incorporation

For BrdU incorporation, embryos were incubated in 2.5 mM BrdU diluted in 1x Embryo Rearing Medium (ERM) for respective amounts of time.

### Induction of Cre/lox system

For ^ERT2^Cre induction, embryos were treated with a 5 µM tamoxifen solution in 1x ERM overnight. For HS::Cre induction, stage 34 embryos were moved to room temperature where the medium was removed completely from the plastic dish and filled with 42°C ERM. Immediately after, the embryos were placed in a 37°C incubator for 2 hours. Finally, they were returned to 28°C.

### Inhibitor treatment

LY-411575 (Sigma Aldrich) was dissolved in DMSO to a 50 mM stock concentration. The stock solution was diluted in 2.5 mM BrdU to reach the final working concentration of 5 μM. Hatching stage fish were treated with 5 μM LY-411575 in BrdU for four days at 28°C in the dark. The solution was exchanged after 2 days.

### Immunohistochemistry on cryosections

Fish were euthanized using Tricaine and fixed overnight in 4% PFA, 1x PTW at 4°C. After fixation samples were washed with 1x PTW and cryoprotected in 30% sucrose in 1x PTW. To improve section quality, the samples were incubated in a half/half mixture of 30% sucrose and Tissue Freezing Medium (Leica) for at least 3 days. 16-µM-thick serial sections were obtained on a Leica cryostat. Sections were rehydrated in 1x PTW for 30 min at room temperature. Blocking was performed for 1–2 hr with 10% NGS (normal goat serum) in 1x PTW at room temperature. The respective primary antibodies were applied diluted 1:500 in 1% NGS o/n at 4˚C. The secondary antibody was diluted 1:750 in 1% NGS together with DAPI (1:500 dilution in 1xPTW of 5 mg/ml stock) and applied for 2h at 37˚C. Slides were mounted with 60% glycerol and kept at 4°C until imaging.

### BrdU immunohistochemistry on cryosections

BrdU antibody staining was performed with an antigen retrieval step. After all antibody stainings and DAPI staining, except for BrdU, were complete, a fixation for 30 min was performed with 4% PFA. Slides were incubated for 1 hr at 37°C in 2 N HCl solution, and pH was recovered by washing with a 40% Borax solution before incubation with the primary BrdU antibody.

### Immunohistochemistry on whole mount retinae

Fish were euthanized using Tricaine and fixed over night in 4% PFA, 1x PTW at 4°C. After fixation samples were washed with 1x PTW. Fish were bleached with 3% H2O2, 0.5% KOH in H2O for 2-3 hours in the dark. Retinae were enucleated and permeabilized with acetone for 15 min at −20°C. Blocking was performed in 1% BSA (Sigma Aldrich), 1% DMSO (Roth/Merck), 4% sheep serum (Sigma Aldrich) in 1xPTW for 2h. Samples were incubated with primary antibody in blocking buffer overnight at 4°C. The secondary antibody was applied together with DAPI in blocking buffer overnight at 4°C. Primary antibodies were diluted 1:200, secondary antibodies 1:250 and DAPI 1:500.

### Antibodies

The following primary antibodies were used: anti-EGFP (chicken; Life Technologies, A10262), rabbit anti-Rx2 (Reinhardt et al., 2015), anti-tagRFP (rabbit, Evrogen, AB233), anti-DsRed (rabbit; Clontech, 632496), anti-BrdU (rat; AbD Serotec, BU1/75), anti-Zpr-1 (mouse; ZIRC). The following secondary antibodies were used (1:500): anti-mouse Cy5 (Jackson ImmunoResearch, 715-175-151), anti-chicken 488 (Jackson ImmunoResearch, 703-485-155), anti-rat DyLight549 (Jackson ImmunoResearch, 112-505-143), anti-rabbit DyLight549 (Jackson ImmunoResearch), anti-mouse Alexa546 (Life Technologies, A-11030) and anti-rat Alexa633 (Life Technologies, A21094). DAPI (Sigma-Aldrich, D9564) nuclear counterstaining was performed as described by Inoue and Wittbrodt (2011).

### Immunohistochemistry imaging

All immunohistochemistry images were acquired by confocal microscopy at a Leica TCS SPE with either a 20x water objective or a 40x oil objective or at a Leica TCS SP8 with 20x or 63X oil objective.

### Image processing and statistical analysis

Images were processed via Fiji image processing software. Statistical analysis and graphical representation of the data were performed using the Prism software package (GraphPad). Unpaired t-tests were performed to determine the statistical significances. The p-value p<0.05 was considered significant and p-values are given in the figure legends. Sample size (n) is mentioned in every figure legend. No statistical methods were used to predetermine sample sizes, but our sample sizes are similar to those generally used in the field. The experimental groups were allocated randomly, and no blinding was done during allocation.

---

**Figure S1. Notch signalling is not active in retinal stem cells in the post-embryonic retina in medaka.** (A) Schematic representation of the retina from a frontal view (0°). (A’) Frontal view of a retina 3D reconstruction. (B) Schematic representation of the retina from a lateral view (90°). (B’) Lateral view of a retina 3D reconstruction. GFP is shown in green and Rx2 in magenta.

**Figure S2. Notch-positive progenitors give rise to Müller glia cells, amacrine cells and bipolar cells.** Endogenous tagRFP signal is shown in A and B (magenta). Immunostaining against tagRFP is shown in C (magenta). A’, B’ and C’ show immunostaining against cell type specific markers (green). A’’, B’’ and C’’ show the merge of tagRFP and the marker. In A’’’, B’’’ and C’’’ the merge together with DAPI nuclear labelling is shown. Scale bar is 20 µm. (A-A’’’) Notch positive progenitors give rise to amacrine cells, detected by immunostaining against Pax6. (B-B’’’) Notch positive progenitors give rise to bipolar cells, detected by immunostaining against PKCα. (C-C’’’) Notch positive progenitors give rise to amacrine cells, detected by immunostaining against Pax6.

**Figure S3. *atoh7*::^ERT2^ Cre induces recombination in progenitor cells of the retina.** (A-A’’) Cryosection of an *atoh7*::^ERT2^ Cre fish crossed to a GaudíRSG fish, induced at hatching and fixed 8 days later. Recombination (green) can be detected in cells located at the beginning of the GCL (arrowhead), directly adjacent to the *rx2* (magenta) expression domain. Few recombined cells are also detected close the beginning of the INL (open arrowhead). Scale bar is 25 μm. (B-B’’) Maximum projection of a whole-mount immunohistochemistry of the retina of *atoh7*::^ERT2^ Cre fish crossed to GaudíRSG fish, induced at hatching and fixed 1 month later. Recombined cells (green) can be detected in a ring around the embryonic retina (asterisk). Scale bar is 200 μm.

**Figure S4. Quantification of the percentage cell types generated from the Atoh7-positive progenitors in each cellular layer in GaudíRSG and GaudíRSNICD.** (A) Quantification of the percentage of GFP-positive (Atoh7-derived) cells shows a decrease in the GCL, ***p=0,0003 (1755 cells in 4 GaudíRSG retinae and 1428 cells in 8 GaudíRSNICD retinae). (B) Quantification of the percentage of GFP-positive (Atoh7-derived) cells shows an increase in the INL, ****p<0,0001 (100 cells in 4 GaudíRSG retinae and 522 cells in 8 GaudíRSNICD retinae). (C) Quantification of the percentage of GFP-positive (Atoh7-derived) cells remained constant in the ONL,^ns^ p=0,2412 (164 cells in 4 GaudíRSG retinae and 268 cells in 8 GaudíRSNICD retinae).

**Figure S5. The inhibitor LY411575 inhibits efficiently Notch signalling.** (A) *tp1::*d2GFP reporter hatch fish were incubated for 4 days in 5 μM LY411575 and then imaged. (B) The reporter fish show fluorescence before the treatment. (C) After 4 days of inhibitor treatment, the fluorescence is clearly reduced and therefore Notch signalling is efficiently inhibited.

## References

Andreazzoli, M. (2009). Molecular regulation of vertebrate retina cell fate. Birth Defects Res. Part C - Embryo Today 87, 284–295.

Atkinson, P. J., Huarcaya Najarro, E., Sayyid, Z. N. and Cheng, A. G. (2015). Sensory hair cell development and regeneration: similarities and differences. Development 142, 1561–1571.

Austin, C. P., Feldman, D. E., Ida, J. a and Cepko, C. L. (1995). Vertebrate retinal ganglion cells are selected from competent progenitors by the action of Notch. Development 121, 3637–50.

Bajoghli, B., Aghaallaei, N., Hess, I., Rode, I., Netuschil, N., Tay, B.-H., Venkatesh, B., Yu, J.-K., Kaltenbach, S. L., Holland, N. D., et al. (2009). Evolution of genetic networks underlying the emergence of thymopoiesis in vertebrates. Cell 138, 186–97.

Baker, N. E., Yu, S. and Han, D. (1996). Evolution of proneural atonal expression during distinct regulatory phases in the developing Drosophila eye. Curr. Biol. 6, 1290–1302.

Bassett, E. A. and Wallace, V. A. (2012). Cell fate determination in the vertebrate retina. Trends Neurosci. 35, 565–73.

Bigas, A. and Espinosa, L. (2012). Hematopoietic stem cells : to be or Notch to be. Blood 119, 3226–35.

Borggrefe, T. and Oswald, F. (2009). The Notch signaling pathway: transcriptional regulation at Notch target genes. Cell. Mol. Life Sci. 66, 1631–46.

Bray, S. J. (2016). Notch signalling in context. Nat. Rev. Mol. Cell Biol. 17, 722–735.

Centanin, L., Hoeckendorf, B. and Wittbrodt, J. (2011). Fate restriction and multipotency in retinal stem cells. Cell Stem Cell 9, 553–62.

Centanin, L., Ander, J.-J., Hoeckendorf, B., Lust, K., Kellner, T., Kraemer, I., Urbany, C., Hasel, E., Harris, W. A., Simons, B. D., et al. (2014). Exclusive multipotency and preferential asymmetric divisions in post-embryonic neural stem cells of the fish retina. Development 141, 3472–3482.

Chitnis, A., Henrique, D., Lewis, J., Ish-Horowicz, D. and Kintner, C. (1995). Primary neurogenesis in Xenopus embryos regulated by a homologue of the Drosophila neurogenic gene Delta. Nature 375, 761–766.

Clark, B. S., Cui, S., Miesfeld, J. B., Klezovitch, O., Vasioukhin, V. and Link, B. A. (2012). Loss of Llgl1 in retinal neuroepithelia reveals links between apical domain size, Notch activity and neurogenesis. Development 139, 1599–1610.

Corson, F., Couturier, L., Rouault, H., Mazouni, K. and Schweisguth, F. (2017). Self-organized Notch dynamics generate stereotyped sensory organ patterns in Drosophila. Science 356, 6337.

Del Bene, F., Wehman, A. M., Link, B. A. and Baier, H. (2008). Regulation of Neurogenesis by Interkinetic Nuclear Migration through an Apical-Basal Notch Gradient. Cell 134, 1055–1065.

Dorsky, R. I., Rapaport, D. H. and Harris, W. A. (1995). Xotch inhibits cell differentiation in the Xenopus retina. Neuron 14, 487–496.

Dorsky, R. I., Chang, W. S., Rapaport, D. H. and Harris, W. A. (1997). Regulation of neuronal diversity in the Xenopus retina by Delta signalling. Nature 385, 67–70.

Easter, S. S. and Johns, P. R. (1977). Growth of the adult goldfish eye, II: Increase in retinal cell number. J. comp Neurol 176, 331–342.

Edlund, T. and Jessell, T. M. (1999). Progression from extrinsic to intrinsic signaling in cell fate specification: A view from the nervous system. Cell 96, 211–224.

Fiúza, U. M. and Arias, A. M. (2007). Cell and molecular biology of Notch. J. Endocrinol. 194, 459–474.

Fre, S., Huyghe, M., Mourikis, P., Robine, S., Louvard, D. and Artavanis-Tsakonas, S. (2005). Notch signals control the fate of immature progenitor cells in the intestine. Nature 435, 964–968.

Graham, T. G. W., Tabei, S. M. A., Dinner, A. R. and Rebay, I. (2010). Modeling bistable cell-fate choices in the Drosophila eye: qualitative and quantitative perspectives. Development 137, 2265–2278.

Hollyfield, J. G. (1968). Differential Addition of Cells to the retina in Rana pipiens Tadpoles. Dev. Biol. 18, 163–179.

Hufnagel, R. B. and Brown, N. L. (2013). Specification of Retinal Cell Types. In Patterning and Cell Type Specification in the Developing CNS and PNS: Comprehensive Developmental Neuroscience, (Chap. 27, Vol. 1, pp. 519–536).

Jensen, J., Pedersen, E. E., Galante, P., Hald, J., Heller, R. S., Ishibashi, M., Kageyama, R., Guillemot, F., Serup, P. and Madsen, O. D. (2000). Control of endodermal endocrine development by Hes-1. Nat. Genet. 24, 36–44.

Jusuf, P. R., Almeida, A. D., Randlett, O., Joubin, K., Poggi, L. and Harris, W. A. (2011). Origin and Determination of Inhibitory Cell Lineages in the Vertebrate Retina. J. Neurosci. 31, 2549–2562.

Jusuf, P. R., Albadri, S., Paolini, A., Currie, P. D., Argenton, F., Higashijima, S., Harris, W. a and Poggi, L. (2012). Biasing amacrine subtypes in the Atoh7 lineage through expression of Barhl2. J. Neurosci. 32, 13929–44.

Kageyama, R., Ohtsuka, T. and Kobayashi, T. (2007). The Hes gene family: repressors and oscillators that orchestrate embryogenesis. Development 134, 1243–1251.

Kay, J. N., Finger-Baier, K. C., Roeser, T., Staub, W. and Baier, H. (2001). Retinal ganglion cell genesis requires lakritz, a Zebrafish atonal Homolog. Neuron 30, 725–736.

Kiernan, A. (2013). Notch signaling during cell fate determination in the inner ear. Semin Cell Dev Biol 24, 479–479.

Kiernan, A. E., Cordes, R., Kopan, R., Gossler, A. and Gridley, T. (2005). The Notch ligands DLL1 and JAG2 act synergistically to regulate hair cell development in the mammalian inner ear. Development 132, 4353–4362.

Kim, D. S., Ross, S. E., Trimarchi, J. M., Aach, J., Greenberg, M. E. and Cepko, C. L. (2008). Identification of molecular markers of bipolar cells in the murine retina. J. Comp. Neurol. 507, 1795–1810.

Koch, U., Lehal, R. and Radtke, F. (2013). Stem cells living with a Notch. Development 140, 689–704.

Lai, E. C. (2004). Notch signaling: control of cell communication and cell fate. Development 131, 965–973.

Link, B. A. and Darland, T. (2001). Genetic analysis of initial and ongoing retinogenesis in the zebrafish: comparing the central neuroepithelium and marginal zone. Prog. Brain Res. 131, 565–577.

Livesey, F. J. and Cepko, C. L. (2001). Vertebrate neural cell-fate determination: lessons from the retina. Nat. Rev. Neurosci. 2, 109–118.

Louvi, A. and Artavanis-Tsakonas, S. (2006). Notch signalling in vertebrate neural development. Nat. Rev. Neurosci. 7, 93–102.

Lust, K., Sinn, R., Pérez Saturnino, A., Centanin, L. and Wittbrodt, J. (2016). De novo neurogenesis by targeted expression of Atoh7 to Müller glia cells. Development 143, 1874–1883.

Matsuda, M. and Chitnis, A. B. (2010). Atoh1a expression must be restricted by Notch signaling for effective morphogenesis of the posterior lateral line primordium in zebrafish. Development 137, 3477–3487.

Maurer, K. A., Riesenberg, A. N. and Brown, N. L. (2014). Notch signaling differentially regulates Atoh7 and Neurog2 in the distal mouse retina. Development 141, 3243–3254.

Merzlyak, E. M., Goedhart, J., Shcherbo, D., Bulina, M. E., Shcheglov, A. S., Fradkov, A. F., Gaintzeva, A., Lukyanov, K. A., Lukyanov, S., Gadella, T. W. J., et al. (2007). Bright monomeric red fluorescent protein with an extended fluorescence lifetime. Nat. Methods 4, 555–557.

Mizutari, K., Fujioka, M., Hosoya, M., Bramhall, N., Okano, H. J., Okano, H. and Edge, A. S. B. (2013). Notch Inhibition Induces Cochlear Hair Cell Regeneration and Recovery of Hearing after Acoustic Trauma. Neuron 77, 58–69.

Neves, J., Abelló, G., Petrovic, J. and Giraldez, F. (2013). Patterning and cell fate in the inner ear: A case for Notch in the chicken embryo. Dev. Growth Differ. 55, 96–112.

Ohnuma, S., Hopper, S., Wang, K. C., Philpott, A. and Harris, W. A. (2002). Co-ordinating retinal histogenesis : early cell cycle exit enhances early cell fate determination in the Xenopus retina. Development 129, 2435–2446.

Parks, A. L., Huppert, S. S. and Muskavitch, M. A. T. (1997). The dynamics of neurogenic signalling underlying bristle development in Drosophila melanogaster. Mech. Dev. 63, 61–74.

Pearson, B. J. and Doe, C. Q. (2004). Specification of Temporal Identity in the Developing Nervous System. Annu. Rev. Cell Dev. Biol. 20, 619–647.

Perron, M. and Harris, W. A. (2000). Determination of vertebrate retinal progenitor cell fate by the Notch pathway and basic helix-loop-helix transcription factors. Cell. Mol. Life Sci. 57, 215–223.

Poggi, L., Vitorino, M., Masai, I. and Harris, W. A. (2005). Influences on neural lineage and mode of division in the zebrafish retina in vivo. J. Cell Biol. 171, 991–999.

Raymond, P. A, Barthel, L. K., Bernardos, R. L. and Perkowski, J. J. (2006). Molecular characterization of retinal stem cells and their niches in adult zebrafish. BMC Dev. Biol. 6, 36.

Reinhardt, R., Centanin, L., Tavhelidse, T., Inoue, D., Wittbrodt, B., Concordet, J., Martinez Morales, J. R. and Wittbrodt, J. (2015). Sox 2, Tlx, Gli 3, and Her 9 converge on Rx 2 to define retinal stem cells in vivo. EMBO J. 34, 1572–1588.

Riesenberg, A. N., Liu, Z., Kopan, R. and Brown, N. L. (2009). Rbpj Cell Autonomous Regulation of Retinal Ganglion Cell and Cone Photoreceptor Fates in the Mouse Retina. J. Neurosci. 29, 12865–12877.

Romero-Carvajal, A., Acedo Navajas, J., Jiang, L., Kozlovskaja-Gumbrien, A., Alexander, R., Li, H. and Piotrowski, T. (2015). Regeneration of sensory hair cells requires localized interactions between the Notch and Wnt pathways. Dev. Cell 34, 267–282.

Schmidt, T. M., Chen, S. K. and Hattar, S. (2011). Intrinsically photosensitive retinal ganglion cells: Many subtypes, diverse functions. Trends Neurosci. 34, 572–580.

Schneider, M. L., Turner, D. L. and Vetter, M. L. (2001). Notch signaling can inhibit Xath5 function in the neural plate and developing retina. Mol. Cell. Neurosci. 18, 458–472.

Siebel, C. and Lendahl, U. (2017). Notch signaling in development, tissue homeostasis, and disease. Physiol. Rev. 97, 1235–1294.

Suzuki, S. C., Bleckert, A., Williams, P. R., Takechi, M., Kawamura, S. and Wong, R. O. L. (2013). Cone photoreceptor types in zebrafish are generated by symmetric terminal divisions of dedicated precursors. Proc. Natl. Acad. Sci. 110, 15109–15114.

Taylor, S. M., Alvarez-Delfin, K., Saade, C. J., Thomas, J. L., Thummel, R., Fadool, J. M. and Hitchcock, P. F. (2015). The bHLH transcription factor neuroD governs photoreceptor genesis and regeneration through delta-notch signaling. Investig. Ophthalmol. Vis. Sci. 56, 7496–7515.

VanDussen, K. L., Carulli, A. J., Keeley, T. M., Patel, S. R., Puthoff, B. J., Magness, S. T., Tran, I. T., Maillard, I., Siebel, C., Kolterud, A., et al. (2012). Notch signaling modulates proliferation and differentiation of intestinal crypt base columnar stem cells. Development 139, 488–497.

Wan, J. and Goldman, D. (2017). Opposing Actions of Fgf8a on Notch Signaling Distinguish Two Muller Glial Cell Populations that Contribute to Retina Growth and Regeneration. Cell Rep. 19, 849–862.

Wan, J., Ramachandran, R. and Goldman, D. (2012). HB-EGF is necessary and sufficient for Müller glia dedifferentiation and retina regeneration. Dev. Cell 22, 334–347.

Wilson, S. G., Wen, W., Pillai-kastoori, L. and Morris, A. C. (2016). Tracking the fate of her4 expressing cells in the regenerating retina using her4 : Kaede zebra fish. Exp. Eye Res. 145, 75–87.

